# Nano-pulling stimulates axon regeneration in dorsal root ganglia by inducing stabilization of axonal microtubules and activation of local translation

**DOI:** 10.1101/2023.10.29.564574

**Authors:** Alessandro Falconieri, Pietro Folino, Lorenzo Da Palmata, Vittoria Raffa

## Abstract

Axonal plasticity is a phenomenon strongly related to neuronal development as well as regeneration. Recently, it has been demonstrated that active mechanical tension, intended as an extrinsic factor, is a valid contribution to the modulation of axonal plasticity. In previous publications, our team validated a method, the “nano-pulling”, for applying mechanical forces on developing axons of isolated primary neurons using magnetic nanoparticles (MNP) actuated by static magnetic fields. This method was found to promote axon growth and synaptic maturation. Here, we explore the possibility to use nano-pulling as an extrinsic factor to promote axon regeneration in a neuronal tissue explant. Having this in mind, whole dorsal root ganglia (DRG) have been dissected from mouse spinal cord, incubated with MNPs, and then stretched. We found that particles were able to penetrate the ganglion and to localise both into the somas and in sprouting axons. Our results point that the nano-pulling doubles the regeneration rate, documented by an increase in the arborizing capacity of axons, and an accumulation of cellular organelles related to mass addition (endoplasmic reticulum and mitochondria) with respect to spontaneous regeneration. In line with the previous results on isolated hippocampal neurons, we observed an increase in the density of stable microtubules and activation of local translation in stretched ganglions. The collected data demonstrate that the nano-pulling is able to enhance axon regeneration in whole spinal ganglia exposed to MNPs and external magnetic fields. The preliminary data represent an encouraging starting point for proposing the nano-pulling as biophysical tool to design novel therapies based on the use of force as an extrinsic factor for promoting nerve regeneration.

## Introduction

Nerve injuries are a serious cause of disability for the entire world population, determining a huge socio-cultural and economic impact. Among the most common causes there are the contusion-related injuries like falls, road and industrial accidents, but also injuries from cuts (e.g., knife, saw blade, fan, glass) or bone fractures (Robinson, 2000; Burnett and Zager, 2004; Campbell, 2008). Nerve compression syndromes, i.e. structural and functional alterations in the nerve or adjacent tissues due to load or pressure, are other typical causes of nerve injuries (Rempel et al., 1999; Burnett and Zager, 2004; Hochman and Zilberfarb, 2004; Campbell, 2008).

To date, a universal method of treating nerve injuries is still missing even if there are various methodologies for their treatment. Unfortunately, there are some hard-to-treat injuries such brain and spinal cord injuries, and most of these diseases are considered uncurable. Recently, the use of nanotechnology in nerve regeneration therapy is gaining increasing interest. The main uses include the production of biomaterials (natural or synthetic) for the development of scaffolds and nerve guidance conduits (Gerth et al., 2015; Sarker et al., 2018) or for implementing smart drug delivery approaches (Tajdaran et al., 2019; Bianchini et al., 2023). Among them, MNPs that have been effectively used for the delivery of growth factors, such as NGF (Ziv-Polat et al., 2014; Zuidema et al., 2015; Giannaccini et al., 2017; Marcus et al., 2018), BDNF (Pilakka-Kanthikeel et al., 2013; Wise et al., 2016), GDNF (Ziv-Polat et al., 2014), VEGF (Giannaccini et al., 2017) to promote nerve survival and neurite regeneration (Zuidema et al., 2015; Wise et al., 2016; Giannaccini et al., 2017). Recently, our team proposed a new application of MNPs for the mechanical stimulation of neurons. Historically, chemical signaling was thought to be the main responsible for neuronal growth and differentiation. However, the role of mechanical force in guiding and promoting these same mechanisms has been now recognized (Suter and Miller, 2011; Franze, 2013; Franze et al., 2013). MNPs have the advantage of exerting extremely low forces, similarly to the endogenous ones (picoNewton order). Since the early years of the 21st century, many groups have exploited MNPs to modulate axonal functions, such as axon specification and orientation (Riggio et al., 2014; Kunze et al., 2015; Raffa et al., 2018), intracellular calcium dynamics (Tay et al., 2016; Tay and Di Carlo, 2017; De Vincentiis et al., 2020), cytoskeletal dynamics (Pita-Thomas et al., 2015; De Vincentiis et al., 2020; Falconieri et al., 2023) axonal transport (Steketee et al., 2011; Chowdary et al., 2013, 2019; Kunze et al., 2017; Falconieri et al., 2023), elongation and branching (Steketee et al., 2011; Riggio et al., 2014; Tay et al., 2016; Raffa et al., 2018; De Vincentiis et al., 2020; Wang et al., 2020; Falconieri et al., 2023), neuron maturation (De Vincentiis et al., 2020, 2023; Falconieri et al., 2023). In our previous studies performed on isolated hippocampal neurons, we found that nano-pulling induces a remodelling of the axonal cytoskeleton, by increasing the number of microtubules (MTs) and the fraction of stable MTs (Falconieri et al., 2023). We suggested that the structural changes at the level of MTs coordinate a modulation of axonal transport, and the activation of local translation, stimulating axon outgrowth and synaptic maturation (Falconieri et al., 2023). These interesting results rise the question whether nano-pulling is relevant for translational research, behind its application as a valuable tool for the study of signal mechanotransduction. In fact, since many MNP-based nano-formulations and magnetic fields are approved for clinical use, nano-pulling has the potential to be used as a non-invasive medical tool/device. However, many points have not been yet confirmed, and questions regarding i) how MNPs interact *in vivo*, ii) whether they are able to penetrate neuronal tissues, iii) whether they can reach the regenerating axons and iv) how they can promote their regeneration still need to be answered. The lack of this knowledge impairs effective translation of the technology in pre-clinical models.

Here, we propose a study on DRGs, bilateral structures located between peripheral nerve terminals and the dorsal horn of the spinal cord, able to carry sensory information (e.g., pain or temperature) from the periphery (peripheral nervous system, PNS) to the central nervous system (CNS) (Berta et al., 2017; Ahimsadasan et al., 2018). DRGs contain not only neurons but also non-neuronal cells (glial cells, endothelial cells, fibroblasts), recapitulating all the complexity of a nerve tissue (Haberberger et al., 2019). Specifically, DRGs are formed by a specific form of glia, called satellite cells, that form an envelope around cell bodies of sensory neurons that project fibers, surrounded by connective tissue and blood vessels (Haberberger et al., 2019). DRG neurons are pseudo-unipolar type of sensory neurons with two branches (the distal process and the proximal process), one projecting into the CNS and the other into the PNS. For this reason, DRGs represent an ideal model system for a pilot study of nerve regeneration, since the dissection maintains the intact structure of the ganglion but the distal process and the proximal processes are resected and their regeneration can be studied under controlled conditions (Nascimento et al., 2018).

Here, this model was used to evaluate the effects of mechanical stimulation mediated by externally-administered MNPs, regardless of the anatomical compartments (CNS *vs* PNS) which, notoriously, present a different predisposition to regeneration (Goldberg and Barres, 2000; Huebner and Strittmatter, 2009; Mietto et al., 2015).

## Methods

### Animals

All the animal procedures were performed in compliance with protocols approved by Italian Ministry of Public Health and of the local Ethical Committee of University of Pisa, in conformity with the Directive 2010/63/EU. Post-natal day (P) 3 C57BL/6J mice were used. They were maintained in a controlled environment (23 ± 1°C, 50 ± 5% humidity) with a 12 / 12 h light / dark cycle with food and water *ad libitum*.

### Dorsal root ganglia organotypic cultures

For DRG organotypic cultures P3 mice were used. For dissection, isolation and culturing we modified a protocol proposed by Han and colleagues (Han et al., 2020). Briefly, animals were sacrificed by decapitation and then, their columns were excised in a dissection medium constituted of a solution of D-glucose 6.5 mg ml^-1^ in DPBS (Gibco; Thermo Fisher Scientific, Waltham, Massachusetts, US; #15050065). The spinal cords were removed and DRGs from cervical and thoracic regions were collected. Then, the DRGs have been stripped of their nerve roots which branch off from the body and placed on glass coverslips previously coated with 500 μg ml^-1^ poly-L-lysine (PLL, Sigma-Aldrich, Burlington, Massachusetts, US, #P4707) and 10 μg ml^-1^ laminin (Gibco, Thermo Fisher Scientific, Waltham, Massachusetts, US, #23017-015). To promote adhesion, the ganglia were placed on ice for 45 minutes. Then, culture medium consisting of Neurobasal-A medium (Gibco, Thermo Fisher Scientific, Waltham, Massachusetts, US, #12348-017) modified with B27 (Gibco, Thermo Fisher Scientific, Waltham, Massachusetts, US, #17504-044), 2 mM Glutamax (Gibco, Thermo Fisher Scientific, Waltham, Massachusetts, US, #35050-038), 50 IU·ml^-1^ penicillin, 50 μg·ml^-1^ streptomycin was added. After four hours, fresh cell culture medium supplemented with 5 μg ml^-1^ MNPs was added, and the samples were incubated at 37°C in a humidity-saturated atmosphere containing 95% air and 5% CO2.

### Magnetic nanoparticles

For magnetic stimulation of DRGs, MNPs were used (Fluid-MAG-ARA, Chemicell, Germany). As stated from the supplier, MNPs were characterized by a core of iron oxide approximately 75±10 nm in diameter, saturation magnetization of 59 Am^2^/kg^-1^ and a hydrodynamic diameter of 100 nm. The outer layer is made of glucuronic acid and represents an organic shell to avoid nanoparticles aggregation.

### Magnetic stimulation

Magnetic stimulation was applied by a Halbach-like cylinder magnetic applicator that provides a constant magnetic field gradient (46.5 T m^-1^) in the radial centrifugal direction (Riggio et al., 2014; Raffa et al., 2018). All the experiments were carried out by placing 35 mm Petri dishes, containing the glass coverslips, inside of the applicator to develop a constant, static and permanent force on the DRGs. Mechanical forces, magnetically-actuated, were applied from Day *in vitro* (DIV) 1 to DIV3.

### Nano-pulling

DRGs (four / five per glass coverslip) were placed in 35 mm Petri dishes at DIV0. After four hours from the complete attachment, MNPs were added (DIV0.17). At DIV1, samples placed inside of the magnetic applicator (stretched groups; Stretch) or outside (control groups; Ctrl). After 48 hours of incubation (DIV3), all the samples were fixed and prepared for fluorescence microscopy.

### Ribopuromycylation

We evaluated the population of ribosomes, actively translating, in DRGs by the ribopuromycylation (RPM) method, modifying a protocol already published (Falconieri et al., 2023) (in turn, modified from (Bastide et al., 2018)). Briefly, DRGs were harvested and seeded on glass coverslips at DIV0. After MNP delivery (DIV0.17) and nano-pulling (from DIV1 to DIV3), the ganglia were treated with 200 μM emetine (Sigma-Aldrich, Burlington, Massachusetts, US, #E2375) and 100 μM puromycin (Sigma-Aldrich, Burlington, Massachusetts, US, #P8833) for 10 min at 37°. Then, ganglia were washed with ice-cold 0.0003% digitonin (Sigma-Aldrich, Burlington, Massachusetts, US; #D141) for 2 min. Next, samples were first washed with ice-cold DPBS and then fixed in 4% PFA, 4% sucrose (Sigma-Aldrich, Burlington, Massachusetts, US, #S0389) for 30 min at RT.

### Immunostaining

At DIV3, DRGs were fixed in 4% PFA and 4% sucrose (Sigma-Aldrich, Burlington, Massachusetts, US, #S0389) for 30 min at room temperature (RT). For studying axon regeneration, samples were permeabilized with 0.5% Triton X-100 in DPBS for 20 minutes and blocked in 5% GS / 0.3% Triton X-100 in DPBS for 45 minutes. BTUBBIII antibody (Sigma-Aldrich, Burlington, Massachusetts, US, #T8578, 1:500) was diluted in 3% GS / 0.2% Triton X-100 in PBS overnight (ON). The day after, samples were washed and incubated with secondary antibody (Thermo Fisher Scientific, Waltham, Massachusetts, US, #06380 or #AB_2633280, 1:500) and Hoechst 33342 (Thermo Fisher Scientific, Waltham, Massachusetts, US, #H3570, 1:1000) for 1 hour at RT.

For staining organelles, we followed a protocol already published (Cioni et al., 2019). Briefly, after fixation, ganglia were removed to study organelle dynamics in the axonal component. Then, samples were permeabilized in 0.1% Triton X-100 in PBS for 5 minutes. Blocking was performed by incubate samples in 5% GS in DPBS for 30 minutes at RT. Primary antibodies (BTUBBIII, Sigma-Aldrich, Burlington, Massachusetts, US, #T8578, 1:500; BTUBBIII, Abcam, Cambridge, UK, #ab41489, 1:1000; acetylated tubulin, Sigma-Aldrich, Massachusetts, US, #T7451, 1:400; tyrosinated tubulin, Abcam, Cambridge, UK, # ab6160, 1:400; KDEL, Thermo Fisher Scientific, Waltham, Massachusetts, US, #PA1-013, 1:200; TOMM20, Abcam, Cambridge, UK, #ab86735; TOMM20, Abcam, Cambridge, UK, #ab289670; Puromycin, Sigma-Aldrich, Burlington, Massachusetts, US, #MABE343, 1:1000; S6, Cell Signaling, Danvers, Massachusetts, US, #2217, 1:200) were diluted in DPBS overnight (ON) at 4°C. The day after, samples were washed and incubated with secondary antibodies (Thermo Fisher Scientific, Waltham, Massachusetts, US, #A11029, #R6393, #A21236, #A11006, #A11008, #A11011, #A21244, #A11041, #A21449, 1:500) and Hoechst 33342 (Thermo Fisher Scientific, Waltham, Massachusetts, US, H1399, 1:1000). Images were acquired using a fluorescent microscope (Nikon, TE2000-U) for nano-pulling assays or a laser scanning confocal microscope (Nikon, Eclipse Ti) for mitochondria, endoplasmic reticulum and ribosomes quantification.

### Samples preparation for transmission electron microscopy

Ultrastructural characterization was carried out in transmission electron microscopy (TEM) following a protocol already published (De Vincentiis et al., 2020). Briefly, DRGs were fixed in 1.5% glutaraldehyde in 0.1 M sodium cacodylate buffer (pH 7.4). After fixation, the ganglions were detached from glass coverslips and the different compartments (ganglions / axons) were processed separately. Then, samples were postfixed in reduced osmium solution (1% OsO4, 1% K3Fe(CN)6, and 0.1 M sodium cacodylate buffer). Staining was performed with our homemade solution (Moscardini et al., 2020). Samples were dehydrated in a growing series of ethanol, and flat-embedded in Epoxy resin (Epoxy embedding medium kit, Merck KGaA, Darmstadt, Germany; #). UC7 LEICA ultramicrotome (UC7, Leica Microsystems) was used to cut ultra-thin sections (90 nm) that were then collected on 300 mesh copper grids (Electron Microscope Science). TEM micrographs were acquired with a TEM microscope JEM-1010 (Jeol, Tokyo, Japan) operating at 80 Kv equipped with MegaView III high-resolution digital camera with an AnalySIS imaging software (Soft Imaging System, Muenster, Germany).

### Image analysis

To establish the different patterns of ramification of DRGs organotypic cultures, the ImageJ software was used (Schneider et al., 2012). Specifically, the branching patterns were evaluated using “neuroanatomy”, a fiji plugin, through the “sholl” function (Ferreira et al., 2014). Specifically, Sholl function was exploited to determine the complessity of ramifications by evaluating the number of intersections of the neurites that arise from the DRGs. Briefly, DRGs were binarized (“threshold” function) and using the “Sholl” function a series of concentric rings from the center of the ganglion were generated. The number of neurites that intersect each ring were counted. The distance between subsequent rings was setted on 5 μm. DRGs axonal ramification was analysed from 4x magnification images. For fluorescence quantification, we evaluated the mean fluorescence 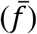, following a method already published (Falconieri et al., 2023). Briefly, the area (A, ROI region of interest), integrated density (IntDen, sum of all the pixel intensities in that selected region) and the mean fluorescence of background readings 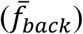 were evaluated in fiji. Specifically, we followed the formula reported here:

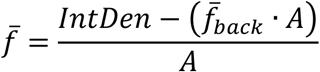

Fluorescence quantification was evaluated from large 10x magnification images taking care to acquire the entire axonal region close to the magnetic field direction.

### Statistical analysis

Data were plotted with GraphPad software, version 7.0.0. Values are reported as the mean ± standard error of the mean (SEM). Outliers were eventually identified and removed using the ROUT method (Q = 1%). The normality of data distribution was tested using the Kolmogorov-Smirnov normality test. We used the *t* test for unpaired data followed by the Bonferroni correction. Mann-Whitney test was performed for non-normally distributed data. Significance was set at *p*≤0.05.

## Results

### Nano-pulling promotes addition of new mass in regenerating axons of DRG explants

DRGs were explanted and the whole ganglions were put in culture (DIV0) and incubated with a MNP-modified medium after a few hours (DIV0.17). We assessed the localization of MNPs by TEM. Figure 1 showed that MNPs localise in the axons of DRG neurons. In particular, TEM micrographs reveal that MNPs can be internalized in the ganglion (fig. S1) and reach the axonal membrane with whose? they establish a strong interaction (fig. 1; green arrows). Considering the absence of cell membrane invaginations in proximity of MNPs, or intracellular vesicles surrounding the MNPs, both active or passive mechanisms of cell internalization could occur. MNPs appear as isolated particles with an inorganic core (iron oxide) and an organic corona (dextran) that serves to prevent aggregation. In fact, as can be seen in figure 1, the MNPs are mainly present as single dots intracellularly. In addition, TEM micrographs show that MNPs within the axoplasm (fig.1; red arrows) are frequently found in the proximity of subcellular compartments such as the endoplasmic reticulum (ER) and mitochondria (m) (cyan arrows).

**Figure 1.**
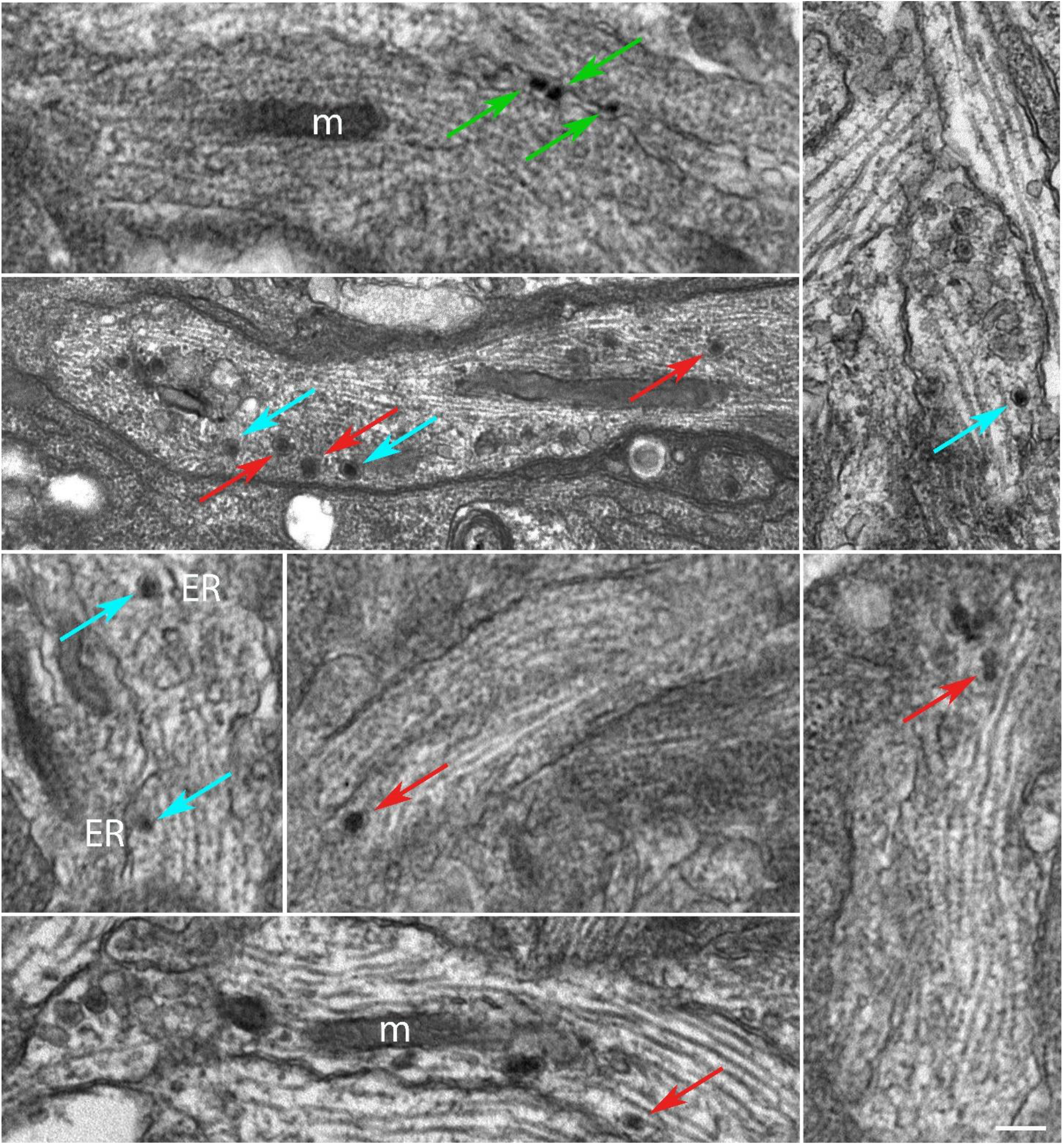
MNPs localize into DRG axoplasm. Red arrows highlight MNPs internalized in the axoplasm of DIV3 DRGs. Green and cyan arrows reveal the strong interactions between MNPs and axonal membrane and sub-cellular components (ER and mitochondria), respectively. Scale bar: 250 nm.

Here, DRG explants were randomly allocated to two groups, the control (Ctrl; fig. 2A1) and the stretched (Stretch; fig. 2A2) groups at DIV1. The stimulation time was set at 48 h, following a procedure that we have already tested in pilot studies on isolated primary neurons and neuron-like cell cultures (Raffa et al., 2018; De Vincentiis et al., 2020). Sholl’s method allowed to verify the effect of mechanical forces on the regenerating axons of DRG explants, by estimating axon branching and ramification. Specifically, we found that the nano-pulling determined a significant increase of the area covered by the axons subjected to stimulation with respect to control ones (fig. 2A3; *p*=0.001). To go deeper into the phenomena of axonal growth in response to tension, we evaluated the elongation rate. Specifically, we found that stretched samples have an elongation rate of 0.025 ± 0.003 μm^2^/h (N= 34 DRG axonal area), i.e., a 2-fold increase compared to spontaneous elongation (0.012 ± 0.001 μm^2^/h; N= 37 DRG axonal area; Mann-Whitney test; *p=*0.003).

**Figure 2.**
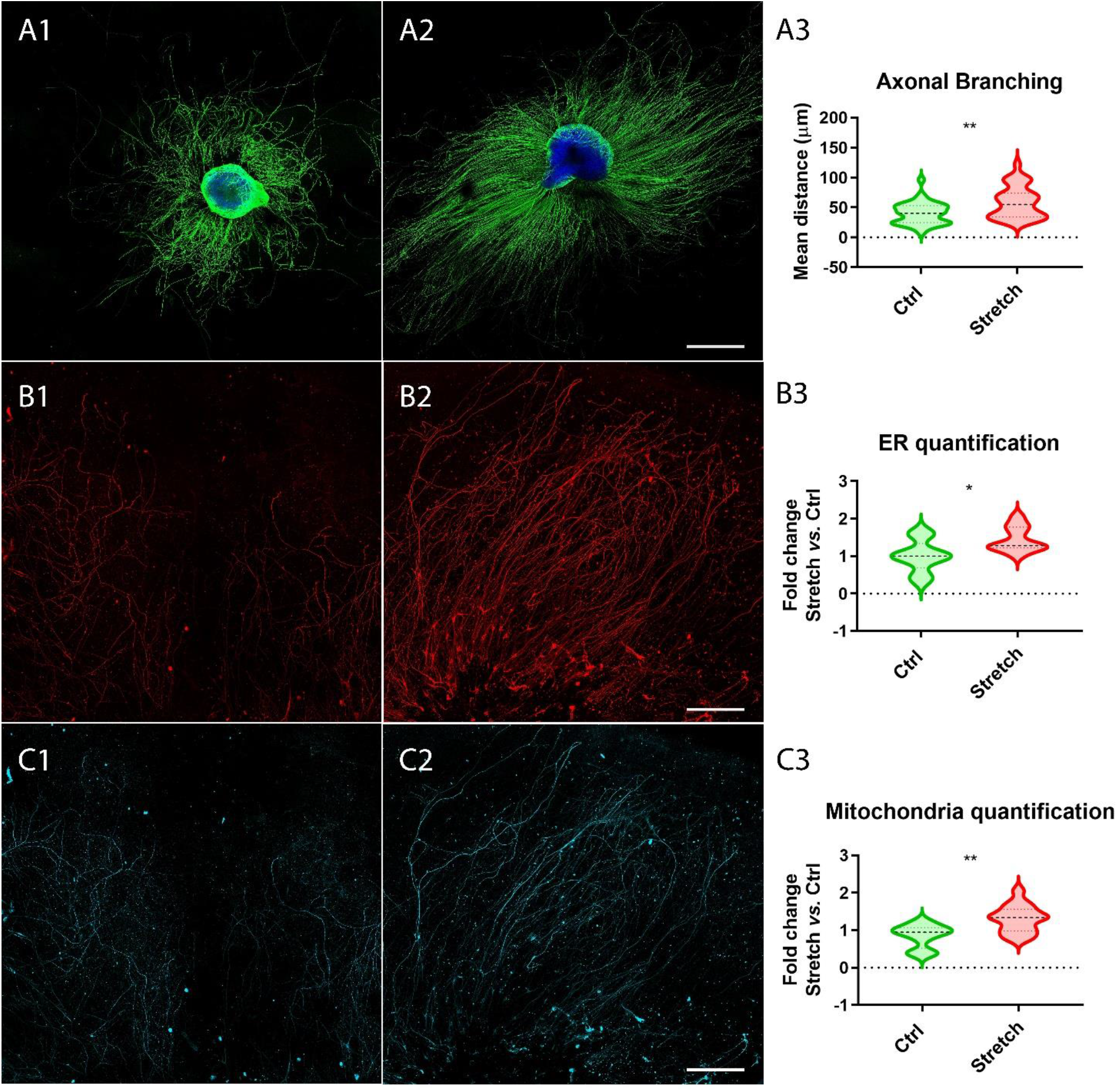
Nano-pulling promotes mass addition in DRG axons. A) DIV3 DRGs in unstretched (A1) and stretched (A2) conditions. B-TUBBIII (green) and DAPI (blue) staining. A3) Sholl analysis of stimulated and unstimulated DRGs. Violin plot (median and extremes as 1^st^ and 3^rd^ quartiles). *t* test for unpaired data. *p*=0.001. N>34 DRGs. B) Immunostaining of endoplasmic reticulum cisternae in unstretched (B1) and stretched (B2) DRGs. KDEL (red) staining. B3) Quantification of ER cisternae. Violin plot (median and extremes as 1^st^ and 3^rd^ quartiles). *t* test for unpaired data. *p*=0.01. N>9 DRGs. C) Immunostaining of mitochondria in control (C1) and stimulated (C2) DRGs. TOMM20 (cyan) staining. C3) Quantification of mitochondria. Violin plot (median and extremes as 1^st^ and 3^rd^ quartiles). *t* test for unpaired data. *p*=0.0016. N>12 DRGs. Scale bars: A = 500 μm; B, C = 250 μm.

Considering that axon growth is usually accompanied by an accumulation of organelles deputed to lipid and protein synthesis and energy supply, we evaluated the amount of endoplasmic reticulum (ER) cisternae (fig. 2B) and mitochondria (fig. 2C) in the regenerating axons sprouting from the ganglion. The whole ganglion was excluded to restrict the analysis to the regenerating component. Experimental data demonstrated that axons subjected to mechanical stimulation (fig. 2B-C2) presented a statistically different amount of ER cisternae (fig. 2B3; *p*=0.01) and mitochondria (2C3; *p*=0.0016) with respect to axons in spontaneous regeneration.

### Nano-pulling promotes activation of local translation in regenerating axons of DRG explants

As local translation is one of the major mechanisms that sustain addition of new mass in the axon, we analysed the fraction of ribosomes in active translation to the total in control samples (fig. 3A1) and in those under nano-pulling (fig. 3A2). We observed a 72% of increase (fig. 3A3; *p*=0.0005) of the ratio of active ribosomes to the total in stretched samples (fig. 3A2, inset) compared to those subjected to spontaneous regeneration (fig. 3A1, inset).

**Figure 3.**
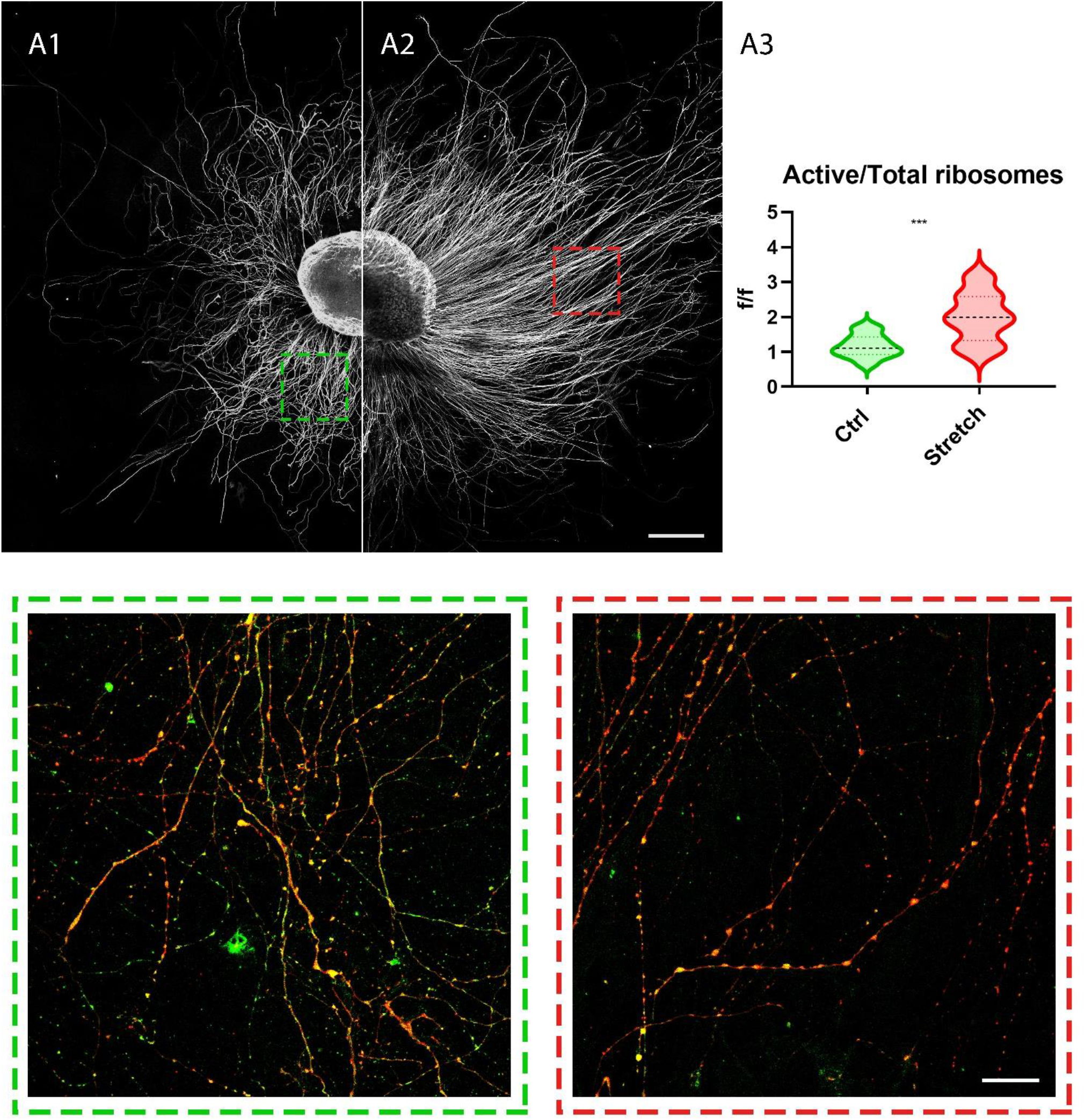
Nano-pulling stimulates the activation of ribosome translation. A) Unstretched (A1) and stretched (A2) DRGs were cultured from DIV1 to DIV3 and the active ribosomes to the total were evaluated in the two conditions (insets) after RPM method. Insets highlight the ratio between active ribosomes and the total population of the same organelles in the condition of spontaneous elongation (A1, inset) and following nano-pulling (A2, inset). Puromycin (red) and S6 (green) staining. B) Analysis of the ratio between active ribosomes to the total in the two conditions. Violin plot (median and extremes as 1^st^ and 3^rd^ quartiles). *t* test for unpaired data. *p*=0.0005. N>12 DRGs. Scale bars: A = 250 μm; Insets = 20 μm.

### Nano-pulling modifies microtubule

One mechanism responsible of activation of local translation is by the assemble of the translational platforms, composed by late endosomes (LE), RNA granules and mitochondria (Cioni et al., 2019). Considering that the transport of the translation machinery is MT-dependent (Broix et al., 2021), we were wondering if the activation of translation could be related to an increase in MT stability, as suggested by our previous studies (Falconieri et al., 2023). To do so, we evaluated the ratio of acetylated to tyrosinated α-tubulin under the two experimental conditions. The rationale behind this is that acetylation is a tubulin post-translational modification that is associated with stability whereas tyrosination is generally related to dynamic instability (Witte et al., 2008). To do this, DRG were seeded on glass coverslips and treated with cell culture medium modified with MNPs after 4 hours for subsequent nano-pulling treatment (DIV0.17). The next day, DRGs were randomly allocated to two groups, control (Ctrl; A1, B1) and nano-pulled (Stretch; A2, B2). After 48 hours of stimulation, we found an increase in the ratio of acetylated to tyrosinated α-tubulin in the shaft of axons subjected to magnetic nano-pulling (fig. 4A3; *p*=0.0013). Conversely, focusing the analysis at the level of the GCs, we found a decrease of acetylated to tyrosinated α-tubulin of stretched samples compared to the condition of spontaneous elongation (fig. 4B3; *p*=0.0016).

**Figure 4.**
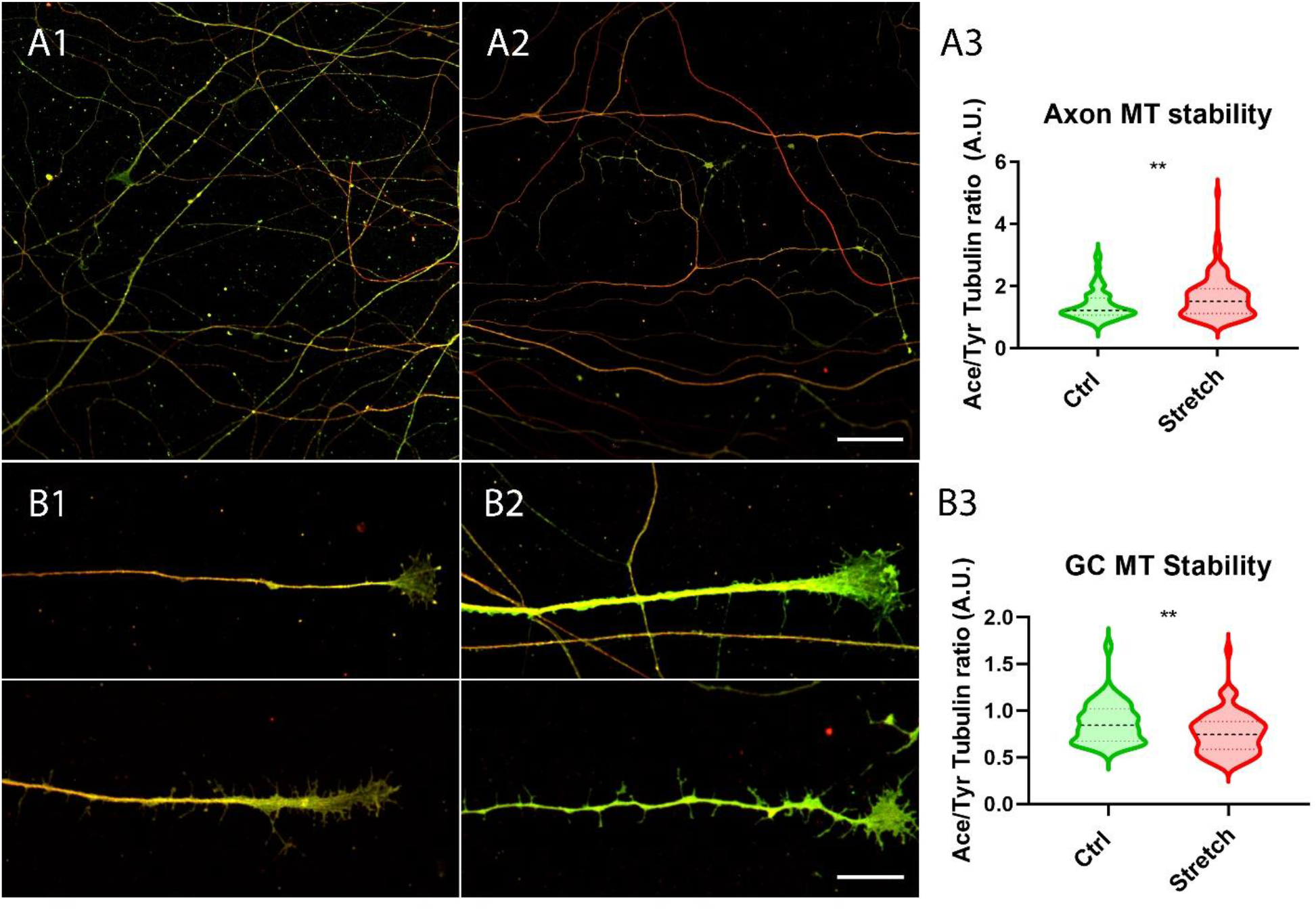
Microtubule stability increases in response to magnetic nano-pulling. A-B) Immunostaining of acetylated (red) *vs* tyrosinated (green) α-tubulin. A) Evaluation of MT stability in control (A1) and stretched (A2) DRG axons. A3) Quantification of the ratio between acetylated and tyrosinated α-tubulin. Violin plot (median and extremes as 1^st^ and 3^rd^ quartiles). Mann-Whitney test. *p*=0.0013. N=125 DRG axons. B) Evaluation of MT stability in unstreched (B1) and stretched (B2) DRG GCs. B3) Quantification of the ratio between acetylated and tyrosinated α-tubulin. Violin plot (median and extremes as 1^st^ and 3^rd^ quartiles). *t* test for unpaired data. *p*=0.0016. N>75 DRG GCs. Scale bars: A = 30 μm; B = 15 μm.

## Discussion

In the present paper, we validated the ability of nano-pulling to induce axon growth in an *in vitro* regeneration model that recapitulates the complexity of a neural tissue, i.e., dissected DRGs. Previous studies shows that nano-pulling is effective only if MNPs are in a close interaction with the axon where they can generate active mechanical force when exposed to a static magnetic field. (De Vincentiis et al., 2020, 2023; Falconieri et al., 2023). Compared to 2D cultures, whole ganglions offer additional barriers to the penetration of MNPs, offering similar features and complexity of *in vivo* neural tissues. MNPs could fail to reach their targets because of the presence of these barriers. First, they could not pass the layers of connective tissue and glial envelope. Second, DRGs contain non-neuronal cells with phagocytic activity (glial cells) that can internalise and quickly destroy the particles. Third, they can be poorly internalised by mature neurons. To validate the ability of MNPs to cross these barriers, soon after the attachment of the ganglions to the glass coverslip, DRGs have been growth in an MNP-modified medium for 3 days. The analysis of the ultrastructure confirms that MNP are able to penetrate the ganglions and to localise inside neural cells (fig. S1). Many particles have been found in the axon, within the axoplasm (red arrows), or interacting closely with the axonal membrane (green arrows). However, the mechanisms of internalisation are difficult to elucidate. Previous works shows that active coatings such as chitosan (Kunze et al., 2017) or wheat germ agglutinin (WGA) (Chowdary et al., 2019) or receptor-specific antibodies (Steketee et al., 2011), promote internalization by endocytosis, as documented by the accumulation of the MNPs in intracellular vesicles. However, our MNPs – that present a glucuronic acid coating - do not show a tendency to cluster in endosome-like vesicles but, rather, they can are found as isolated particles floating in the cytoplasm, attached to organelles or interacting with the cell membranes (Fig. 1). Direct penetration could be an alternative mechanism to endocytosis as it has been shown that some nanoparticles can be internalised directly (Zhang et al., 2011) but deeper studies are needed to confirm this hypothesis. For the scopes of the present study, it was crucial to confirm the presence of MNPs in DGR neurons and a very strong interaction between MNPs and one or more axonal components (e.g. membrane, ER and mitochondria), supporting the assumption that they could induce force generation when manipulated with magnetic fields. This is corroborated by our experimental finding that demonstrated that the nano-pulling increase the axon regeneration rate of about a 2-fold factor. We were wondering which is the molecular component or mechanisms that might be involved in evoking these modulations. Here, we found that the ratio of acetylated versus tyrosinated α-tubulin is decreased in the growth cone of stretched axons, documenting the presence of dynamic MTs (Baas and Black, 1990)that is a typical feature of elongating axons. In fact, rapidly growing GCs are characterised by the presence of highly dynamic, exploratory MTs that translocate from the central domain to the transition domain of the GC during tip advance (Lee and Suter, 2008; Schaefer et al., 2008; Athamneh et al., 2017).

Conversely, this ratio was found to increase in the shaft of stretched axons, documenting the presence of more stable MTs (Takemura et al., 1992; Cappelletti et al., 2021). In general, an increase in acetylation of MTs, with a consequent increase in stability, has been always observed in various studies on neurons (Morley et al., 2016; Yan et al., 2018; Teoh et al., 2022), as well as in other work on non-neuronal models (Swiatlowska et al., 2020; Coleman et al., 2021). We found in previous works that the stabilization of the MTs in the shaft causes their accumulation in the axon (De Vincentiis et al., 2020, 2023; Falconieri et al., 2023). This mechanism seems to be independent on the technology used for force generation, as a similar trend was also observed with magnetic microposts, a different magnetically-activated technology, but capable of exerting higher forces extracellularly (Falconieri et al., 2022).

We previously demonstrated that the most direct consequence of the accumulation of MT in the axon shaft is the positive modulation of the MT-dependent transport of vesicles and organelles. Among organelles, mitochondria and endoplasmic reticulum are fundamental to sustain the production of proteins/lipids necessary for growth and energy to support this production during regeneration. In energy production, a key role is played by the mitochondria, which, at the cellular level, provide for the production of adenosine triphosphate (ATP), which when hydrolysed into diphosphate (ADP) releases energy required for a variety of cellular functions. Thus, we have shown that nano-pulling results in an increase in the amount of mitochondria compared to the control condition (fig. 2C). This data on mitochondria in stretched ganglia are in line with previous finding on mice, human and chick isolated neurons (Lamoureux et al., 2010; De Vincentiis et al., 2023; Falconieri et al., 2023)

Endoplasmic reticulum is another key component for the production of proteins required for axon regeneration. Our experimental data revealed a strong increase in ER cisternae in stretched samples compared to spontaneous regeneration (fig. 2B). The accumulation of ER cisternae as an effect of nano-pulling had already been observed in previous studies on isolated mouse hippocampal neurons (De Vincentiis et al., 2020; Falconieri et al., 2023) and human neural stem cells (De Vincentiis et al., 2023).

Generation of force with technologies other than nano-pulling has the same effect on ER and mitochondria accumulation (Falconieri et al., 2022).

The third component required for in situ production of proteins are ribosomes involved in local translation. Data obtained showed that nano-pulling resulted in a strong increase of active ribosomes in stretched axons (fig. 3). In the previous study carried out by our team, we observed that local translational also occurs through the formation of translation platforms between late endosomes and both active ribosomes and RNA granules (Falconieri et al., 2023). The first group to theorise and demonstrate the formation of these platforms for local translation was Holt’s group in 2019, in whose work they discussed functional contacts between the components involved in the translation machinery for the production of newly synthesised proteins (Cioni et al., 2019). Mitochondria, along with late endosomes and RNA granules, serve as a key component of these platforms. The increase in the concentration of mitochondria, ER, and active ribosomes observed in the present work strongly supports the hypothesis of the formation of platforms useful for local translation and protein production during nano-pulling.

In conclusion, we recently proposed a model according to which the mechanical stimulation induced (by an unknown mechanisms) stabilization of axonal MTs, resulting in accumulation of MT and MT-dependent transport of organelles and vesicle, which, in turn, favour the assembly of the “translational platform” and activation of local translation. Here, we extend the validity of this model to another case study, i.e. a model of axon regeneration represented by dissected whole DRGs.

The data obtained in this work represent an encouraging starting point for the application of nano-pulling technology for promoting the regeneration of a neural tissue. The use of MNPs in biomedicine is widespread and the idea of exploiting them to promote neuroregeneration is gaining great interest (Falconieri et al., 2019). This study on DRGs showed that it is also possible to exploit nano-pulling at the tissue level. However, although DRGs are more informative than 2D cultures, an organotypic model could not recapitulates the complexity of an *in vivo* system. The necessary future direction will be to understand whether this technology can also be used within a living organism by promoting regeneration in a damaged or diseased neural tissue.

## Acknowledgements

The study was supported by the Wings for Life Foundation (WFL-IT-16/17 and 20/21), the EC programme Horizon2020 (101007629-NESTOR-H2020-MSCA-RISE-2020), the Human Frontier Science Program (RGP0026/2021), and the European Union Next-GenerationEU - National Recovery and Resilience Plan (NRRP) – mission 4 component 2, investiment n. 1.4 – CUP N. B83C22003930001 (Tuscany Health Ecosystem – THE”, Spoke 8). This manuscript reflects only the authors’ views and opinions, neither the European Union nor the European Commission can be considered responsible for them. The authors thank Dr. Valentina Cappello and Allegra Coppini for the preparation of TEM samples and Dr. Patrizia Nardini and Daniele Guasti for TEM analysis. We acknowledge the Imaging Platform, Department of Experimental & Clinical Medicine, section Anatomy & Histology of the University of Florence for the access to the TEM facility.

