## Supplemental figure 1 for "Nano-pulling stimulates axon regeneration in dorsal root ganglia by inducing stabilization of axonal microtubules and activation of local translation"

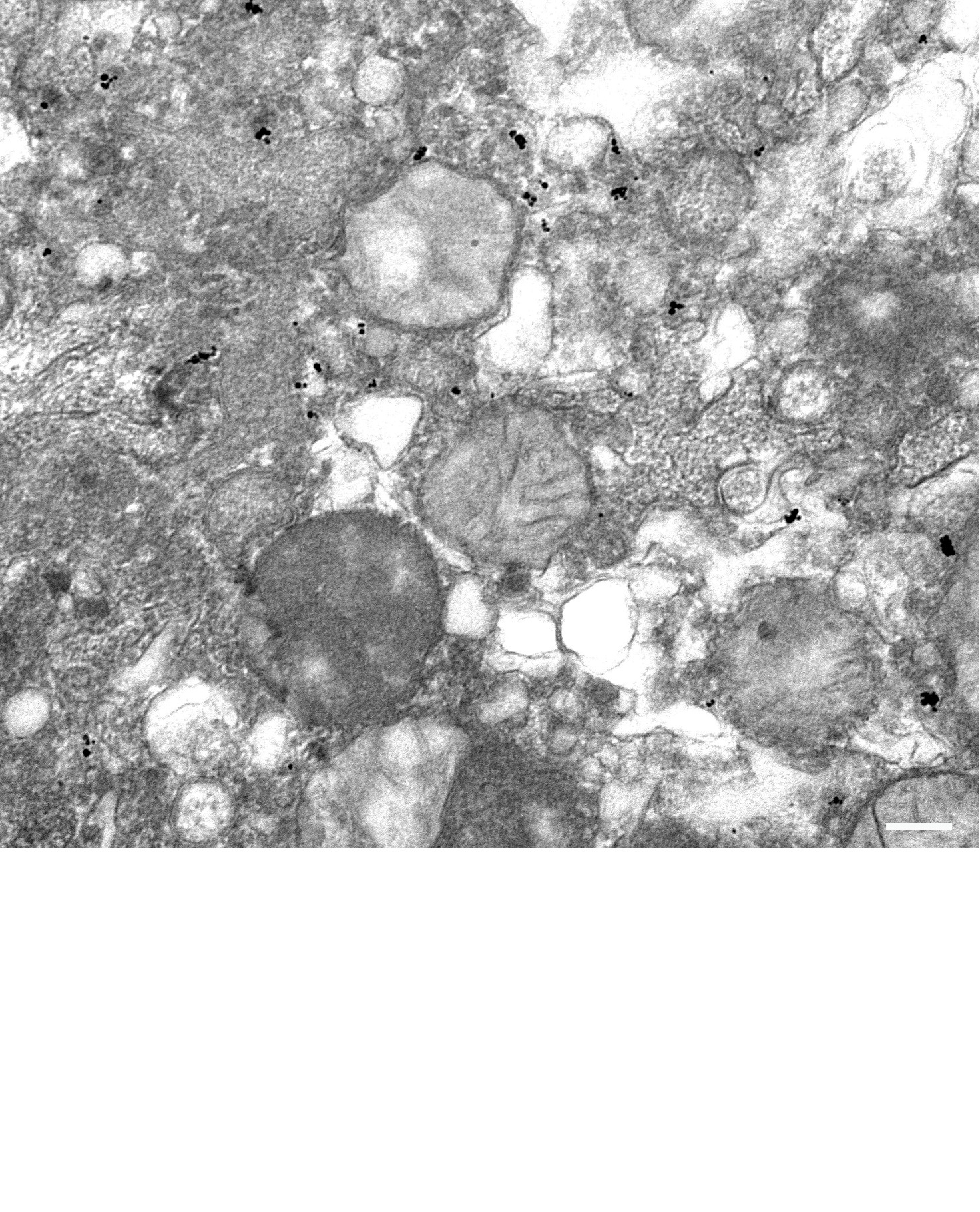
**Figure S1.** *MNPs are internalized into DRGs.* MNPs are mainly present as single dots but, sometimes, they form little clusters of particles differently to axoplasm in which the majority are individual particles. Scale bar: 150 nm.
